# Melatonin attenuates mitochondrial and metabolic dysfunction caused by leptin deficiency

**DOI:** 10.1101/466995

**Authors:** Yaiza Potes, Andrea Díaz-Luis, Juan C Bermejo-Millo, Zulema Pérez-Martínez, Beatriz de Luxán-Delgado, Adrian Rubio-González, Ivan Menéndez-Valle, José Gutiérrez-Rodríguez, Juan J Solano, Beatriz Caballero, Ignacio Vega-Naredo, Ana Coto-Montes

**Affiliations:** Department of Morphology and Cell Biology, Faculty of Medicine, University of Oviedo, 33006, Oviedo, Spain; Instituto de Investigación Sanitaria del Principado de Asturias (IISPA), 33011 Oviedo, Spain; Microbiology service, University Central Hospital of Asturias, 33011, Oviedo, Spain; Geriatric Service, Monte Naranco Hospital, 33012, Oviedo, Asturias, Spain

**Keywords:** Leptin, obesity, metabolism, mitochondria, melatonin

## Abstract

Leptin, as a nutritional inhibitor by repressing food intake, is critical compromised in the major common forms of obesity. Skeletal muscle is the main effector tissue for energy expenditure modifications by the effect of endocrine axes, such as leptin signaling. Our study has been carried out using skeletal muscle from leptin-deficient animal model, in order to ascertain the importance of this hormone in eating disorders. Here we report that leptin-deficiency stimulates an uncontrolled oxidative phosphorylation metabolism, resulting in an excess of energy production that culminates in mitochondrial dysfunction. Thus, different nutrient sensing pathways are perturbed, loosing proteostasis and promoting lipid anabolism, that induces myofiber degeneration and drives oxidative type I fiber conversion. Melatonin treatment plays a significant role in regulating energy homeostasis and fuel utilization. This study reveals melatonin to be a decisive mitochondrial function-fate regulator, with implications for resembling physiological energy requirements and targeting glycolytic type II fibers recovery.

## Introduction

The emerging epidemic of obesity in developed countries is a major risk factor for growing markedly obesity-related comorbidities including cardiovascular diseases, musculoskeletal disorders and certain cancers, resulting in increased mortality (World Heath Organization, 2015). Physiological decline of leptin levels or leptin resistance results in hyperphagic behavior and nutrient overload, which are the central features of eating disorders. Skeletal muscle has an efficient capability of nutrient uptake and storage and, due to this, is considered an important metabolic modulator of glucose, amino acid and lipid homeostasis (Lipina & Hundal, 2017). Thus, leptin directly regulates glucose and fatty acids metabolism in skeletal muscle fibers (Ceddia, 2005). Then, leptin-signaling disruption which also alters circadian rhythms of metabolic activity and feeding (Szewczyk-Golec, Wozniak et al., 2015), could be pivotal for skeletal muscle damage and dysfunction.

Early obesity can alter muscle stem cells by reducing their function and number and impairing their ability to differentiate, ultimately leading to muscle mass loss (Sinha, Sakthivel et al., 2017, Woo, Isganaitis et al., 2011). In that case and given the postmitotic nature of skeletal muscle cells, quality control mechanisms seem to be determinant for the maintenance of cellular homeostasis and skeletal muscle function in obese individuals. Particularly, quality control autophagy, which is mainly regulated by the nutrient-sensing mTOR pathway, supports bioenergetics demands and contributes to metabolic adaptation during cell fate determination (Tang & Rando, 2014). Likewise, mitochondrial bioenergetics could be a linker between leptin disorders and muscle loss or maintenance. Lack of leptin causes an energy imbalance due the high dietary energy intake and the suppression of energy expenditure (Szewczyk-Golec et al., 2015). Simultaneously, nutrient overload induces an intensive oxygen metabolism leading to an abnormal mitochondrial production of reactive oxygen species (ROS) that exceeds the requirements of normal physiological responses (Wellen & Thompson, 2010). Thus, alterations in mitochondrial function and structure could be considered as primary instigators of muscle atrophy under nutrient overload. Mitochondrial function and biogenesis are important factors operating on muscle mass regulation, as well as, on fiber type transition (Liu, Liang et al., 2016, Romanello & Sandri, 2015). Therefore, mitochondrial bioenergetics and quality control mechanisms in skeletal muscle fibers are considered as potential targets for obesity-related muscle atrophy.

To understand the role of leptin in skeletal muscle fibers bioenergetics and quality control, we examined here the skeletal muscle from leptin-deficient obese mice. Furthermore, as melatonin has recognized antioxidant and chronobiotic properties (Chakir, Dumont et al., 2015, Sharafati-Chaleshtori, Shirzad et al., 2017), one can ask whether melatonin treatment could mimic leptin effects on skeletal muscle fibers from leptin-deficient mice. Here, we show that leptin-deficiency reprograms energy requirements of muscle fibers, driving a boost in energy demand largely covered by mitochondria to such an extent that culminates in an impairment of oxidative phosphorylation and adaptive capacity of mitochondria. In addition, this altered interplay between leptin signaling and energy homeostasis perturbs quality control mechanisms leading to the loss of proteostasis in skeletal muscle fibers and to the promotion of lipid anabolism that ultimately induce fast-to-slow muscle fiber type conversion and skeletal muscle dysfunction and degeneration.

## Results

### Leptin deficiency alters skeletal muscle mass

Leptin-deficient ob/ob mice gained double the weight from baseline to sacrifice as the group of control mice, but reduced muscle weight by 27%. Ob/ob mice also showed a higher fat free mass index (FFMI) that was associated with a greater body mass index (BMI). The skeletal muscle index (SMI), FFMI and limb-appendicular skeletal muscle mass index (L-ASMI) are common widely used and have been proposed to study body distribution changes (Zamboni, Mazzali et al., 2008). Interestingly, ob/ob mice experimented an increase in FFMI and a decrease in SMI and L-ASMI, which indicates the coexistence of higher levels of fat mass, particularly intramuscular and peripheral fat, with a musculoskeletal mass decrease in leptin-deficient mice (Table EV1). As well as, the color of skeletal muscle shifted from dark red in wild-type mice to pale red in ob/ob mice, which could be due to the increase in fat infiltration (Fig EV1). Melatonin treatment did not affect these parameters. Thus, leptin deficiency produces a direct impact on skeletal muscle maintenance and composition, which makes the study of bioenergetics and its implication in skeletal muscle quality control mechanisms of paramount importance.

### Skeletal muscle from leptin-deficient obese mice presents altered mitochondrial function, oxidative stress, inflammation and lipid homeostasis

To investigate the implications of the absence of leptin signaling on mitochondrial bioenergetics, we compared the levels of some subunits of the mitochondrial electron transport chain (ETC) complexes, namely NADH dehydrogenase (ubiquinone) 1 β subcomplex 8 (NDUFB8) from complex I; succinate dehydrogenase (ubiquinone) iron-sulfur subunit (SDHB) from complex II; ubiquinol-cytochrome c reductase core protein II (UQCRC2) from complex III; cytochrome c oxidase subunit I (MTCO1) from complex IV and ATP synthase subunit α (ATP5A) from complex V by immunodetection. Thus, we found that leptin-deficient mice present alterations in the ETC machinery, characterized by a reduced expression of subunits from complexes I, IV and V and increased levels of SDHB. In fact, the expression of the mitochondrial mass marker porin was higher in skeletal fibers from ob/ob mice indicating higher amounts of mitochondria than in wild-type mice. The administration of melatonin to ob/ob mice increased the protein levels of NDUFB8 and SDHB subunits and greatly reduced porin expression (Fig 1A). To evaluate if this remodeling of the ETC impacts on mitochondrial function, we studied oxygen consumption and ATP production. Mitochondrial respiration rates were determined in isolated mitochondria from skeletal muscles using a Clark-type oxygen electrode. According to data obtained from state 3 respiration using glutamate/malate as respiration substrates, there were no differences in the ADP-stimulated respiration between the four experimental groups. Despite this, when analyzing state 4 respiration (respiration in the absence of ATP synthesis), we found that ob/ob mice show a higher oxygen consumption than wild-type mice, suggesting that the activity of the ETC is higher in ob/ob mice. The treatment with melatonin did not induce any change in state 4 respiration in both genotypes. The ratio between state 3 and 4 (RCR) indicated no differences in the coupling between substrate oxidation and oxidative phosphorylation in the presence of glutamate/malate. However, the effectiveness of oxidative phosphorylation (ADP/O) in the presence of glutamate/malate was lower in ob/ob animals, an effect reverted by the treatment with melatonin (Fig 1B). By contrast, when we used succinate as substrate for ETC with rotenone and therefore, without taking into account respiration from complex I, state 4 and ADP/O were largely preserved in ob/ob mice. Hence, respiration from complex II in obese animals maintained the effectiveness of oxidative phosphorylation in spite of a slight decrease in RCR that is likely to be connected to proton leak rise and respiration uncoupling. Furthermore, in this case melatonin was also able to increase ADP/O in ob/ob animals (Fig 1C). Despite these alterations in mitochondrial respiration, mitochondria from ob/ob mice maintained their membrane potential that was increased by the treatment with melatonin (Fig EV2). Paradoxically, skeletal muscles from ob/ob mice showed significantly higher mitochondrial and cytosolic ATP levels than those from wild-type mice. Melatonin was able to restore mitochondrial ATP levels and reduce cytosolic ATP levels (Fig 1D). Altogether, our results suggest a higher mitochondrial function in ob/ob mice. The lack in leptin-mediated satiety probably increases cellular energy demand increasing respiratory fluxes to produce higher amounts of ATP. However, we found higher state 4 respiration rates and reduced RCR values in ob/ob animals that would indicate the occurrence of non-respiratory oxygen consumption, associated with the conversion of oxygen to superoxide radical. Accordingly, we found higher lipid peroxidation levels in muscle fibers from ob/ob mice that was accompanied by an imbalanced antioxidant defense with low cytosolic superoxide dismutase (SOD) activity and high cytosolic catalase (CAT) and glutathione peroxidase (GSH-Px) activities (Fig 2A and 2B). Moreover, mitochondrial SOD and CAT activities were lower in leptin-deficient mice (Fig 2C). In addition, this high oxidative stress in ob/ob mice was accompanied by increased levels of interleukin 6 (IL-6) and tumor necrosis factor alpha (TNF-α) (Fig EV3). Melatonin administration was able to reduce lipid peroxidation rates by increasing cytosolic SOD, CAT, glutathione reductase (GR) and mitochondrial CAT activities, while inflammation markers remained unaltered. To further investigate the relationship between oxidative stress and mitochondrial function in skeletal muscle from ob/ob mice, we also analyzed the p66 isoform of SHC (Src homologous-collagen homologue adaptor) protein which presents a dual role as a nutritional sensor and mitochondrial pro-oxidant adaptor (Berniakovich, Trinei et al., 2008). Extracellular stress and mitochondrial dysfunction induces the phosphorylation of p66SHC that is then translocated to mitochondria where enhances further mitochondrial ROS production. Leptin-deficient mice exhibited no alterations in the other SHC isoforms (p46SHC and p52SHC) and an activation of the oxidative stress-dependent p66SHC pathway. Despite melatonin administration increased the total levels of p66SHC in ob/ob mice, the activated p66SHC pathway persisted unaltered (Fig 2D).

**Figure 1.**
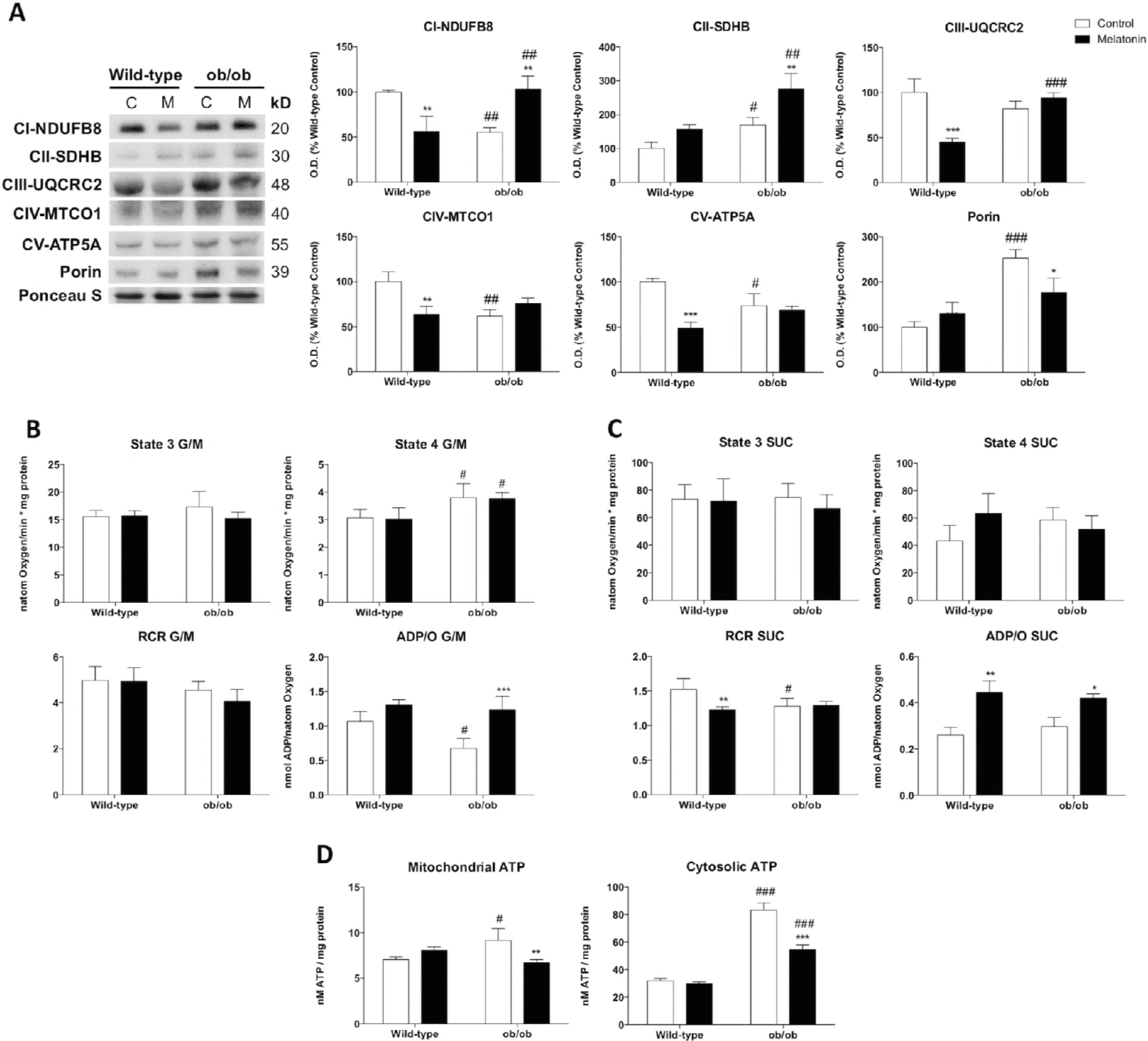
Leptin-deficiency remodels OXPHOS complexes and alters mitochondrial respiration. (A) Protein expression analysis of OXPHOS subunits from each complex (NDUFB8, SDHB, UQCRC2, MTCO1 and ATP5A) and porin. Data are mean of optical density (O.D.) ± SD expressed as percentage of wild-type control mice. Porin and ponceau staining were used as loading control. (B and C) ADP-stimulated O_2_ consumption (State 3), respiration in the absence of ADP-stimulation (state 4), respiratory control ratio (RCR) and ADP/O were analyzed using glutamate/malate (G/M) and succinate (SUC) as respiration substrates of complexes 1 and 2 respectively. Data are mean ± SD (D) Basal mitochondrial and cytosolic ATP content. Data are mean ± SD Statistical comparisons: ^#^ wild-type vs. ob/ob; ^*^Control vs. Melatonin. The number of symbols marks the level of statistical significance: one for *P* < 0.05, two for *P* < 0.01 and three for *P* < 0.001

**Figure 2.**
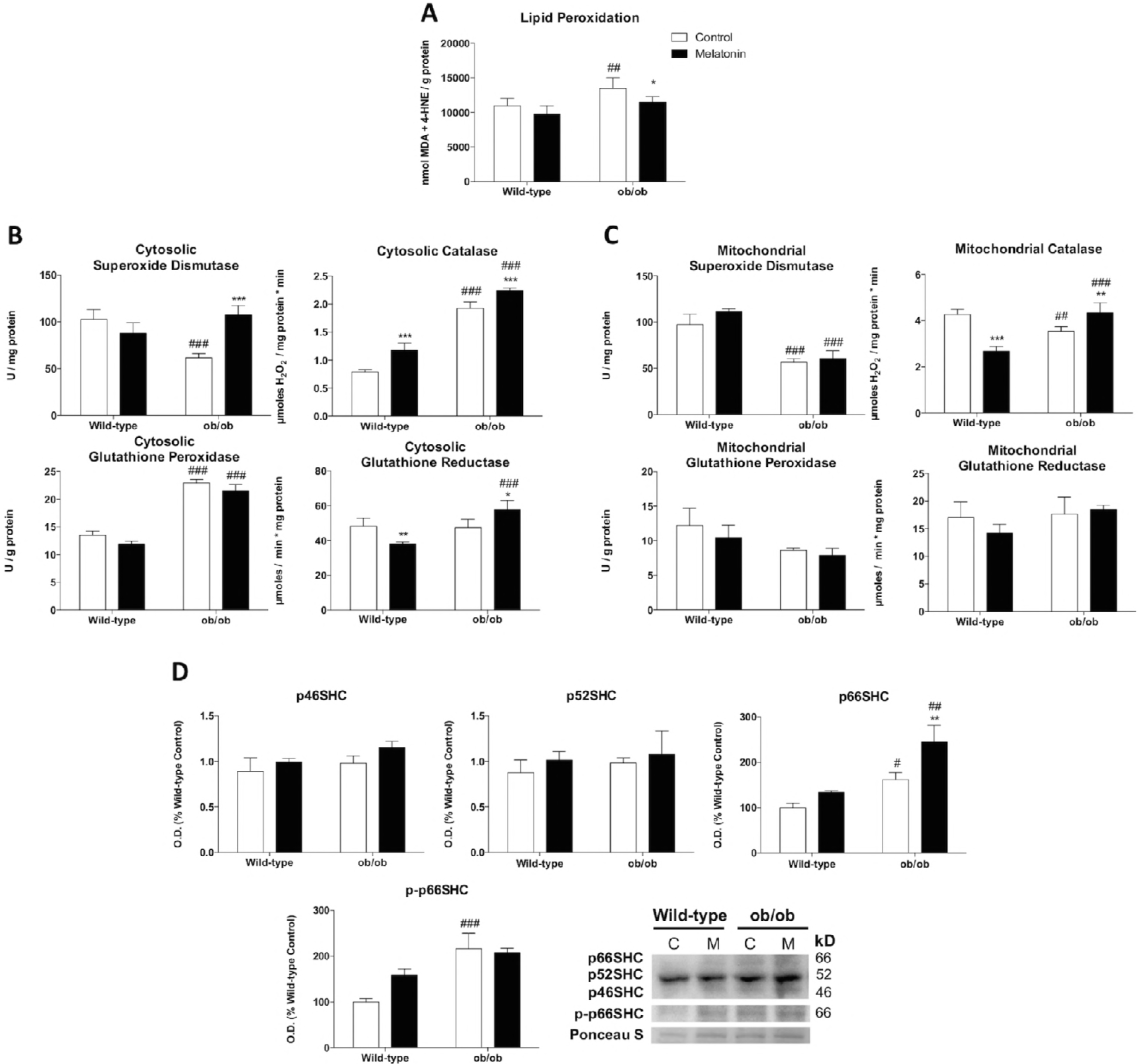
Leptin-deficiency induces skeletal muscle damage and alters mitochondrial and cytosolic antioxidant defense. (A) Determination of lipid oxidative damage (lipid peroxidation). Data are mean ± SD. (B and C) Antioxidant system evaluation by the activity determination of antioxidant enzymes (superoxide dismutase, catalase, glutathione peroxidase and glutathione reductase) in cytosolic and mitochondrial fractions. Data are mean ± SD. (D) Protein expression analysis of Shc-transforming protein 1 (p46Shc, p52Shc, p66Shc and pp66Shc). Data are mean of optical density (O.D.) ± SD expressed as percentage of wild-type control mice. Ponceau staining was used as loading control. Statistical comparisons: ^#^ wild-type vs. ob/ob; ^*^Control vs. Melatonin. The number of symbols marks the level of statistical significance: one for *P* < 0.05, two for *P* < 0.01 and three for *P* < 0.001

Our results would situate skeletal muscle mitochondria from ob/ob mice in a scenario of energy overload that may lead to mitochondrial stress and dysfunction. In addition, we found a downregulation in lactate dehydrogenase protein expression (Fig EV4A) and higher levels of mitochondria-bound hexokinase II (Fig EV4B) in leptin deficient obese mice suggesting anaerobic metabolism inhibition, a higher coupling between glycolysis and oxidative phosphorylation and the inhibition of mitochondrial apoptosis. In fact, muscle mitochondria from ob/ob mice showed lower levels of Bcl-2-associated X protein (BAX) and apoptosis-inducing factor (AIF) accompanied by increased mitochondrial cytochrome c expression indicating a lower susceptibility to events triggering mitochondrial outer membrane permeabilization (Fig EV4C). In addition, we also detected, in ob/ob mice, an overexpression of cyclophilin D, a regulatory component of the mPTP (Giorgio, von Stockum et al., 2013) that exerts apoptosis-suppressing effects mediated by mitochondrial binding of hexokinase-II (Machida, Ohta et al., 2006) (Fig EV4D). Despite the reduction in mitochondrial-bound hexokinase-II and in cyclophilin D expression induced by melatonin treatment in ob/ob mice, mitochondrial BAX and AIF levels remained unaltered. However, mitochondrial cytochrome c levels were reduced by melatonin action.

Mitochondrial energy metabolism is determinant for the maintenance of lipid homeostasis. Similarly, the transcriptional activation of peroxisome proliferator-activated receptor alpha (*Ppara*) is considered as a nutritional sensor being fundamentally important for energy homeostasis and fatty acid metabolism in tissues with high oxidative rates such as muscle (Pawlak, Lefebvre et al., 2015, Wang, 2010). Thus, to evaluate the effect of the absence of leptin on mitochondrial β-oxidation and fatty acid synthesis in skeletal muscle from ob/ob mice, we analyzed PPAR-α signaling. Despite mRNA levels of adiponectin receptors 1 (*Adipor1*) and 2 (*Adipor2*) showed no major changes, the satiety response attenuated by leptin deficiency increased *Ppara* mRNA expression that initially suggests an enhanced mitochondrial β-oxidative capacity, while maintaining the mRNA expression levels of peroxisome proliferator-activated receptor gamma (*Pparg*), a master regulator of adipogenesis. Conversely, higher mRNA expression of acetyl-CoA carboxylase (*Acaca*) was found in obese mice suggesting the promotion of fatty acid synthesis pathway and the inhibition of fatty acid β-oxidation (Fig 5A). To further investigate if *Ppara* altered-response could drive an incomplete fatty acid oxidation and the development of insulin resistance and glucose intolerance (Finck, Bernal-Mizrachi et al., 2005, Koves, Ussher et al., 2008), the c-Jun N-terminal kinase (*Jnk*) cascade was also analyzed. Although mRNA levels of insulin receptor (*Insr*) experimented a significant increase, the absence of leptin also triggered *Jnk* expression in muscle fibers (Fig EV5B). Thus, leptin deficiency might lead to lipid mitochondrial overload by incomplete β-oxidation contributing to alterations in insulin and glucose homeostasis. Despite the fact that melatonin treatment led to higher *Acaca* expression, was also able to increase *Adipor2* and *Pparg* levels in muscle fibers from ob/ob mice.

### Leptin deficiency stimulates mitochondrial biogenesis and fusion

To further investigate whether leptin deficiency-induced obesity may underlie other mitochondrial adaptations in skeletal muscle fibers, we studied mitochondrial biogenesis and dynamics. Leptin-deficient mice showed higher mRNA levels of peroxisomal proliferator-activated receptor coactivator 1 alpha (*Ppargc1a*) and mitochondrial transcription factor A (*Tfam*), suggesting the induction of mitochondrial biogenesis (Fig 3A). Moreover, under stressful conditions and metabolic cues, mitochondrial morphology rapidly changes to optimize mitochondrial bioenergetics. Our results revealed important changes in the molecular machinery of mitochondrial dynamics showing a depletion of fission protein dynamin-related protein 1 (DRP1) and the upregulation of the fusion protein mitofusin 1 (MFN1) in ob/ob animals. However, mitofusin 2 (MFN2) remained unaltered (Fig 3B). Altogether, these results seem to indicate the accumulation of fused mitochondria in skeletal muscle fibers from ob/ob mice. Melatonin treatment increased the expression of *Ppargc1a* in leptin-deficient mice, without changing *Tfam* levels, probably strengthened *Ppargc1a* action as a regulator of ROS metabolism. Moreover, melatonin showed no effects on DRP1 content in ob/ob animals while downregulated MFN1 levels but upregulated MFN2 protein which is involved in the restoration of mitochondrial tubules (Chen, Detmer et al., 2003). Thus, melatonin seems to alter fusion-related mechanisms contributing to the improvement of mitochondrial function.

**Figure 3.**
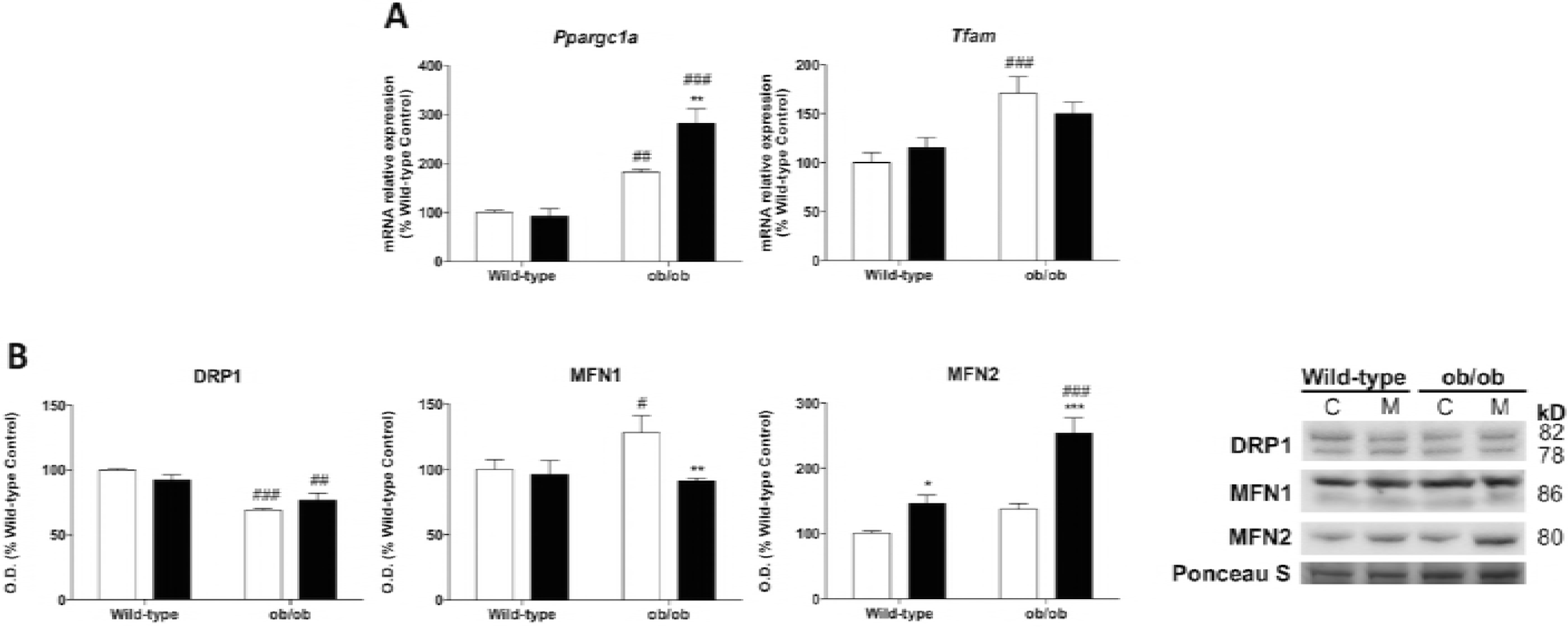
Leptin-deficiency triggers mitochondrial remodeling pathways. (A) Relative mRNA expression of mitochondrial biogenesis genes (*Tfam* and *Ppargc1a*). Data are mean of mRNA relative expression ± SD expressed as percentage of wild-type control mice. Ponceau staining was used as loading control in protein analysis. (B) Protein expression analysis of mitochondrial remodeling markers (DRP1, MFN1 and MFN2). Data are mean of optical density (O.D.) ± SD expressed as percentage of wild-type control mice. Ponceau staining was used as loading control. Statistical comparisons: ^#^ wild-type vs. ob/ob; ^*^Control vs. Melatonin. The number of symbols marks the level of statistical significance: one for *P* < 0.05, two for *P* < 0.01 and three for *P* < 0.001

### Leptin deficiency compromises UPR activation

At this moment we demonstrated disturbances in energy homeostasis that lead to a metabolic reprogramming in skeletal muscle cells from ob/ob animals that would activate additional adaptive responses. Since a major stress response that is activated under metabolic stress signals is the unfolded protein response (UPR) (van der Harg, van Heest et al., 2017), additional analyses were conducted to determine if the three UPR arms were activated. Obese ob/ob mice exhibited reduced inositol-requiring enzyme 1 alpha (IRE1α) protein levels and increased X-box binding protein 1 (*Xbp1*) mRNA expression. The functionally active transcription factor XBP1s resulted from the spliced *Xbp1* mRNA also showed a decline (Fig 4A). Altogether, these results indicate that IRE1α/XBP1 pathway that is the point of confluence of endoplasmic reticulum (ER) stress, insulin signaling and inflammation (Hu, Han et al., 2006, Urano, Wang et al., 2000), was deactivated in leptin-deficient mice. Activating transcription factor 6 alpha (ATF6α) pathway was increased in ob/ob animals (Fig 4B) indicating the activation of adaptive responses against the stress induced by the accumulation of unfolded/misfolded proteins in the ER of muscle fibers from ob/ob mice. However, the phosphorylation at Ser51 of the translation initiation factor alpha (eIF2α) pathway that is involved in the reduction of global protein synthesis (Dever, 1999), exhibited a deactivation in ob/ob mice. Despite this, C/EBP homologous protein (CHOP), a key mediator of the ER stress-mediated apoptosis pathway showed no differences between groups (Fig 4C). Therefore, skeletal muscle cells from obese ob/ob mice seem to display alterations in protein homeostasis and/or deregulated UPR adaptive responses. Melatonin treatment produced a deactivation of the ATF6α pathway and an even greater deactivation of IRE1α/XBP1 and eIF2α pathways.

**Figure 4.**
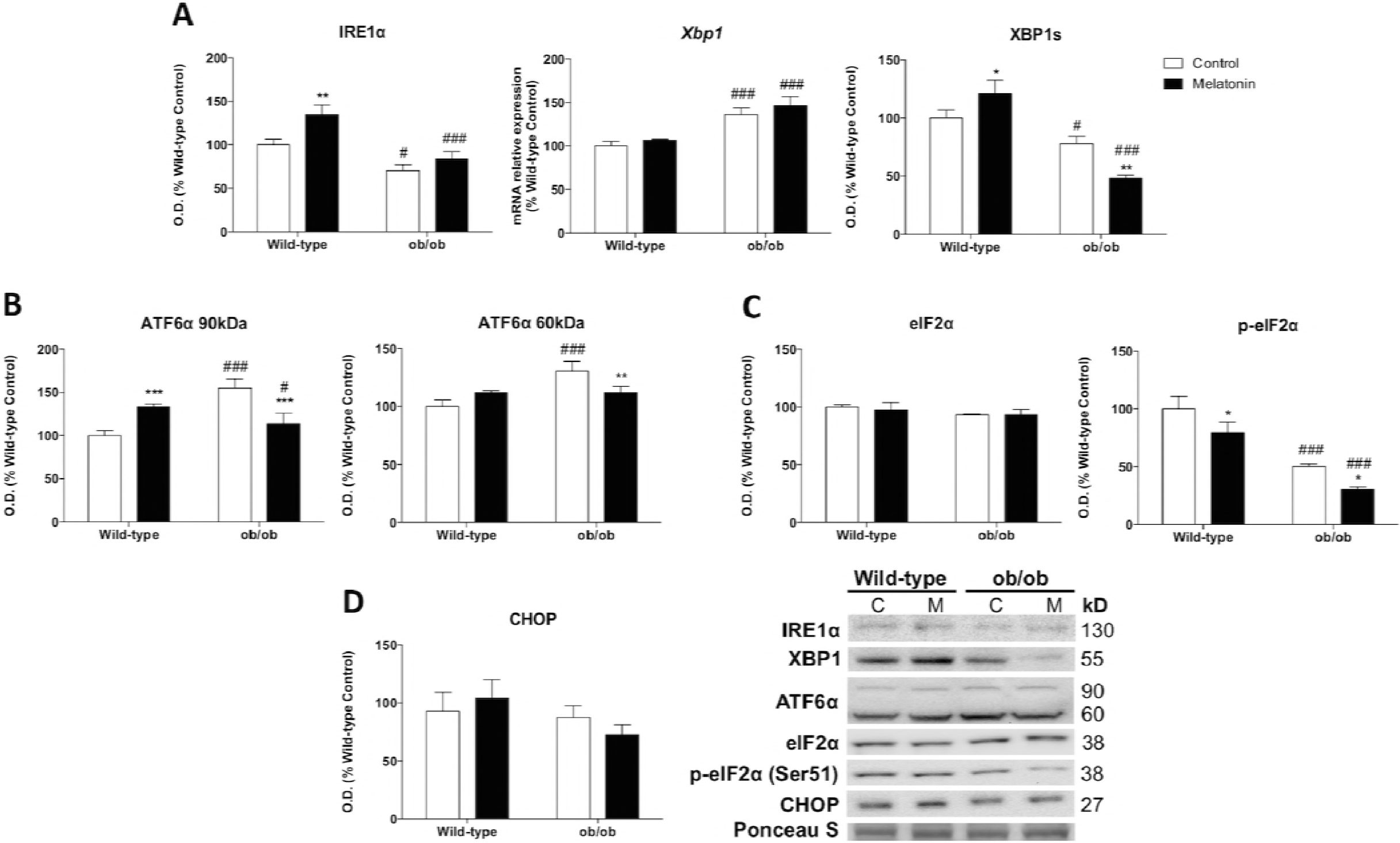
Leptin-deficiency alters UPR. (A) Protein and relative mRNA expression of markers involved the UPR branch activated by IRE1 α and the specific splicing of *Xbp1* (IRE1α, *Xbp1* and XBP1s). Data are mean of optical density (O.D.) or mRNA relative expression ± SD expressed as percentage of wild-type control mice. Ponceau staining was used as loading control. (B) UPR signaling branch activated by a specific proteolytic cleavage of ATF6α was analyzed. Data are mean of optical density (O.D.) ± SD expressed as percentage of wild-type control mice. Ponceau staining was used as loading control. (C and D) UPR signaling markers from branch activated by eIF2α phosphorylation (eIF2α, p eIF2α and CHOP) were analyzed. Data are mean of optical density (O.D.) ± SD expressed as percentage of wild-type control mice. Ponceau staining was used as loading control. Statistical comparisons: ^#^ wild-type vs. ob/ob; ^*^Control vs. Melatonin. The number of symbols marks the level of statistical significance: one for *P* < 0.05, two for *P* < 0.01 and three for *P* < 0.001

### Leptin deficiency inhibits protein translation and enhances autophagy

We then further explore how deficiency of leptin in skeletal muscle fibers affected proteostasis. Ob/ob mice showed no major changes in phosphoinositide-3-kinase (PI3K) protein expression and a lower phosphorylated protein expression of protein kinase B (AKT) and mammalian target of rapamycin (mTOR) at Ser473 and Ser2448, respectively (Fig 5A). In addition to this down-regulation of the AKT/mTOR pathway, the ribosomal protein S6 kinase (p70S6K) phosphorylated at Thr389 was also reduced (Fig 5B), indicating the suppression of this survival pathway and probably increasing protein turnover and degradation to ameliorate ER stress. Then, we also evaluated the activation of the lysosomal-autophagy protein degradation system. We found increased Beclin-1 and microtubule-associated protein 1 light chain 3 (LC3) converted to autophagosome-associating form (LC3-II) protein levels and decreased content of the ubiquitin-binding protein p62 (p62) in ob/ob animals, indicating a higher autophagic flux (Fig 5C). In addition, chaperone-mediated autophagy (CMA), determined by lysosome-associated membrane protein type 2a (*Lamp2a*) mRNA levels, was also triggered in muscle fibers from obese mice (Fig 5D). Induction of autophagy by melatonin treatment in ob/ob mice was unnecessary since Beclin-1 and LC3-II expression were reduced maintaining low levels of p62. However, CMA response was triggered by melatonin action in comparison to not treated obese animals.

**Figure 5.**
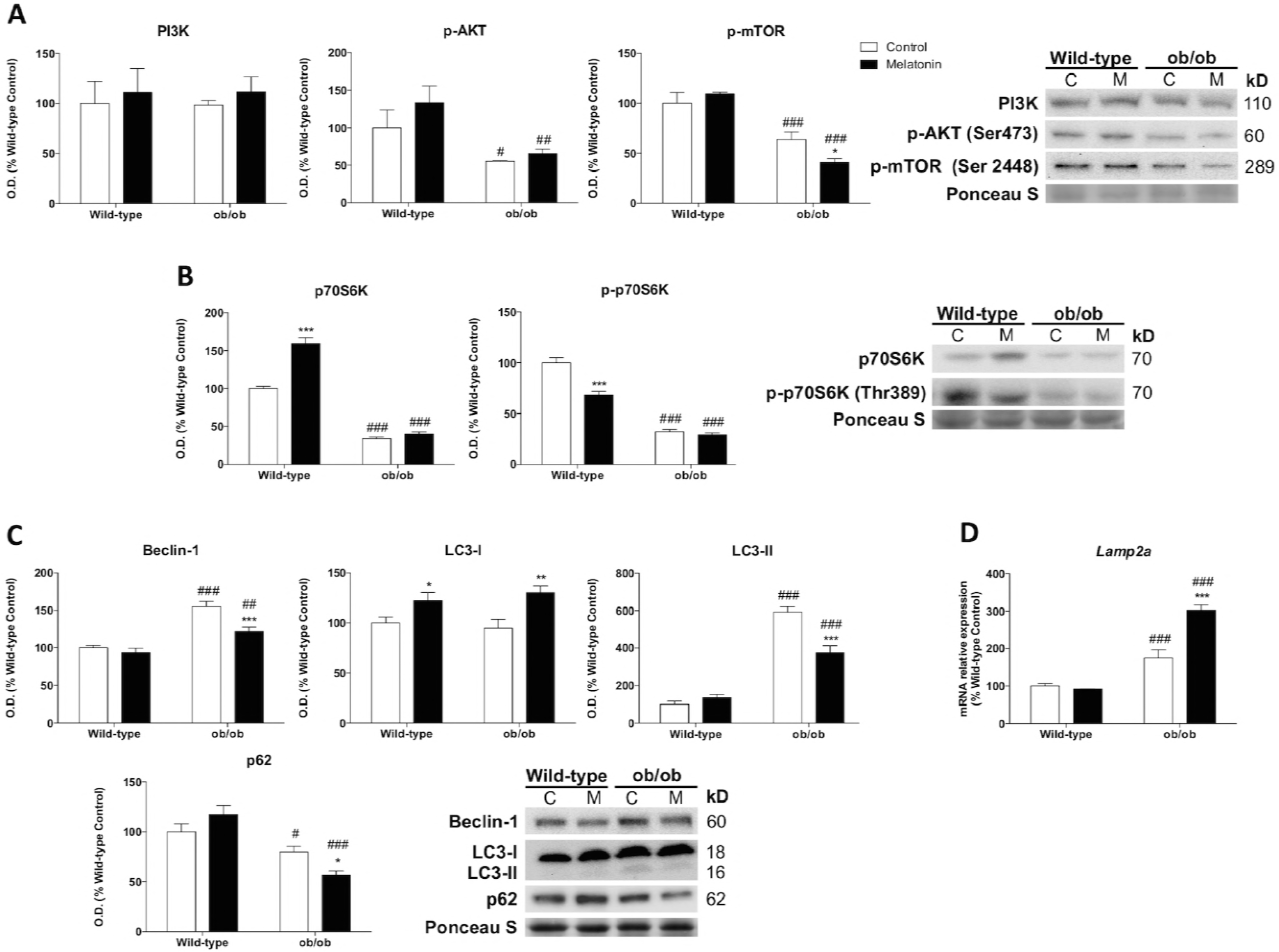
Leptin-deficiency reduces protein synthesis and increases autophagy response via mTOR inhibition. (A) PI3K, p-AKT and p-mTOR protein content show significant changes between genotype remodeling protein synthesis and autophagy responses. Data are mean of optical density (O.D.) ± SD expressed as percentage of wild-type control mice. Ponceau staining was used as loading control. (B) Protein expression analysis of markers downstream of protein biosynthesis pathway (p70S6K and p-p70S6K). Data are mean of optical density (O.D.) ± SD expressed as percentage of wild-type control mice. Ponceau staining was used as loading control. (C) Protein levels of autophagy mechanism (Beclin-1, LC3-I, LC3-II and p62). Data are mean of optical density (O.D.) ± SD expressed as percentage of wild-type control mice. Ponceau staining was used as loading control. (D) Relative mRNA expression of LAMP2A implicated in chaperone-mediated autophagy analysis to evaluate protein removal upon mild-oxidative stress. Data are mean ± SD expressed as percentage of wild-type control mice. Statistical comparisons: ^#^ wild-type vs. ob/ob; ^*^Control vs. Melatonin. The number of symbols marks the level of statistical significance: one for *P* < 0.05, two for *P* < 0.01 and three for *P* < 0.001

### Leptin-deficiency drives glycolytic conversion to oxidative myofiber type and disturbs muscle excitation-contraction coupling

The metabolic reprograming described here in ob/ob mice was also accompanied by a remodeling in skeletal muscle tissue. We found a functional muscle fiber-type switching toward the formation of oxidative type I fibers. Quantitative studies of periodic acid-Shiff (PAS) muscle stained sections showed that obese animals presented a smaller percentage of glycolytic type II fibers, indicating that the absence of leptin favors oxidative fibers. Consistently, the protein expression of muscle ring-finger protein-1 (MURF1), that regulates type II fiber trophicity and maintenance (Moriscot, Baptista et al., 2010), was reduced in ob/ob mice confirming the loss of glycolytic type II fibers (Fig 6A). Furthermore, skeletal muscle of leptin-deficient mice showed a lack of activation of ryanodine receptor 1 (RYR1) channel via phosphorylation. P(Ser2808)-RYR1 depletion accompanied by decreased phosphorylated Ca^2+^/calmodulin-dependent protein kinase II (CaMKII) at Thr286 indicate the inhibition of calcium release by sarcoplasmic reticulum (Fig 6B). Melatonin treatment was able to restore the percentage of type II fibers and to increase P(Ser2808)-RYR1, CaMKII and P(Thr286)-CaMKII levels slightly improving the excitation-contraction coupling.

**Figure 6.**
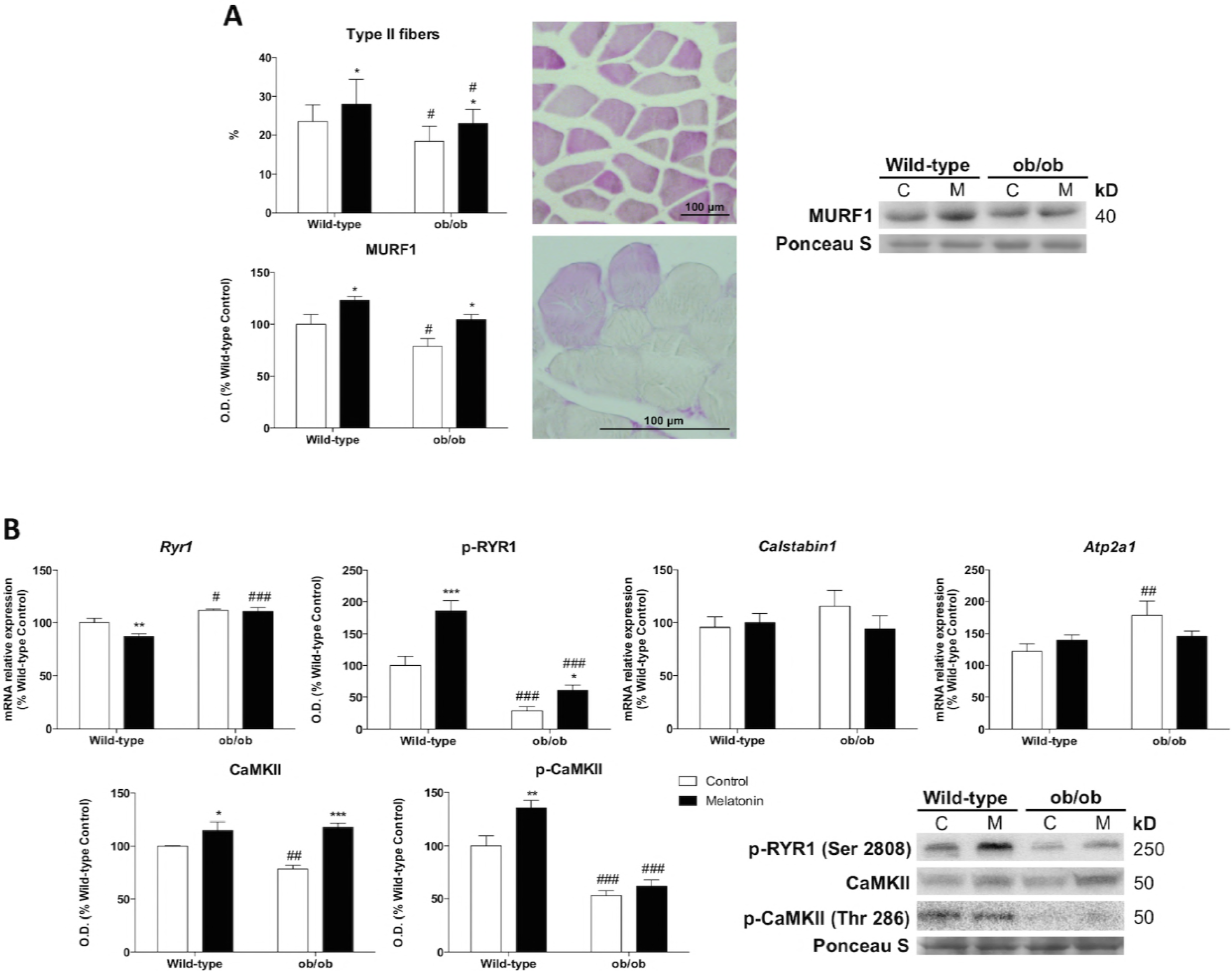
Leptin-deficiency alters fiber type proportion and diminishes muscle excitation-contraction coupling. (A) Light microscopy images of periodic acid-Schiff (PAS) stained sections from skeletal muscle tissue and type II fibers darker stained proportion determination. Scale bars: 100 μm. Protein expression of MURF1 related to increased proportion of type II fibers.Data from protein expression analysis are mean of optical density (O.D.) ± SD expressed as percentage of wild-type control mice. Ponceau staining was used as loading control. (B) Relative mRNA and protein expression of markers involved in skeletal muscle excitatition-contraction pathway (*Ryr1*, p-RYR1, *Calstabin1*, *Atp2a1*, CaMKII and p-CaMKII). Data are mean of mRNA relative expression or optical density (O.D.) ± SD expressed as percentage of wild-type control mice. Ponceau staining was used as loading control. Statistical comparisons: ^#^ wild-type vs. ob/ob; ^*^Control vs. Melatonin. The number of symbols marks the level of statistical significance: one for *P* < 0.05, two for *P* < 0.01 and three for *P* < 0.001

## Discussion

Leptin-deficiency, due to the constant feeling of hunger, leads to an excessive calorie intake that results in pronounced fat mass accumulation. However, the role of leptin seems to be also essential for skeletal muscle growth and maintenance (Hamrick, Herberg et al., 2010) and in fact, our results revealed a severe skeletal muscle loss in ob/ob mice. However, the primary cellular mechanisms underlying the impaired leptin signaling-associated skeletal muscle wasting remain unknown. In the present work, we have demonstrated that leptin deficiency induces a metabolic reprogramming in the skeletal muscle fibers that acquire an energy conservation profile that compromises muscle mass and functional performance. It was described that in a situation of energy deficiency, less leptin is secreted (Bluher, Shah et al., 2009). Thus, the absence of leptin in ob/ob mice provides a constant signal indicating a state of limited energy supply, which induces an enormous energy demand, ultimately compromising mitochondrial physiology. Obese mice force skeletal muscle cells to rely more on OXPHOS for ATP production, compromising its effectiveness from complex I. However, complex II, which acts as a compensatory process when complex I capacity is defective (Hawkins, Levin et al., 2010), is able to support mitochondrial metabolism, meeting the great ATP production demanded under leptin deficiency. Despite this, the low RCR values observed in ob/ob animals indicate that their mitochondria have a low capacity for substrate oxidation and ATP turnover (Brand & Nicholls, 2011). Thus, the lack of leptin simultaneously induces high ATP production by mitochondrial respiration stimulation, as well as a high occurrence of non-respiratory O_2_ consumption that leads to ROS production. In accordance with our results, increased ROS and the activation of antioxidant defense have been previously described in disuse muscle atrophy (Pellegrino, Desaphy et al., 2011). Here, we have also observed a self-triggered loop of ROS production in mitochondria from ob/ob mice via the activation of the pro-oxidant p66SHC-signaling pathway and the poor mitochondrial antioxidant capability. Defective mitochondrial function is correlated with a higher phosphorylation status of p66SHC and vice versa (Lebiedzinska, Karkucinska-Wieckowska et al., 2010, Nemoto, Combs et al., 2006). Therefore, our findings suggest that p66SHC could be a crucial linker between nutrient and energy availability and mitochondrial function.

Overexpression of mitochondria-bound hexokinase-II also plays an important role in the metabolic remodeling elicited by leptin deficiency, thereby contributing to a high glycolytic profile. This is of particular importance in rapidly growing cells such as cancer cells (Mathupala, Ko et al., 2006), or non-tumor pluripotent cells (Varum, Rodrigues et al., 2011), that promotes the association of hexokinase-II and the anaerobic production of lactate to get quick energy boost. In obesity, in spite of the great importance of energy metabolism status, this connection remained unknown. Our results show that skeletal muscle from leptin-deficient mice switch to a higher mitochondria-hexokinase-II affinity due the high-energy demand. However, lactic acid fermentation is critically compromised, thereby, glycolysis is coupled to mitochondrial metabolism that is facilitated by mitochondrially bound hexokinase-II (Roberts & Miyamoto, 2015). It was described that this mitochondria-hexokinase-II association is also able to inhibit apoptosis by interfering with the ability of Bax to bind to mitochondria, inhibiting mitochondrial outer membrane permeabilization (Giorgio et al., 2013, Pastorino, Shulga et al., 2002). Indeed, leptin deficiency reduced Bax translocation to the mitochondria and increased levels of cyclophilin D, leading to the inhibition of mitochondrial release of cytochrome c and AIF. Despite not being released to the cytosol, mitochondrial AIF levels in obese mice were extremely low, an effect that could be relevant for mitochondrial functioning in ob/ob mice. It was described that AIF knockdown disrupts complex I function increasing superoxide production in mitochondria (Varecha, Paclova et al., 2012). Then, our data seems to support a mitochondrial function impairment thorough complex I and an increase in mitochondrial ROS production in skeletal muscle fibers from ob/ob mice. Our results also demonstrate that leptin-deficient mice suppress apoptosis that could be triggered as an adaptive response. In fact, apoptotic cell death contributes to the maintenance of cellular homeostasis and is required for skeletal muscle differentiation (Fernando, Kelly et al., 2002). Therefore, apoptosis inhibition in skeletal muscle from ob/ob mice may have long-term cumulative effects being detrimental for cell proliferation and skeletal muscle repair. This reprogramming of energy metabolism also comprises lipid metabolism. A scenario of energy overload is of paramount importance for mitochondria that serves as the principal lipid sensors of skeletal muscle (Koves et al., 2008). However, the constant exposure to an excessive energy demand induced by the lack of leptin enhances *Ppara* signaling, that is activated under conditions of energy deprivation, promoting fatty acids catabolism (Desvergne & Wahli, 1999). However, a recent study has demonstrated that obesity induces a switch in mitochondrial fuel that relies on carbohydrate substrate utilization, leading to mitochondrial lipid overload and incomplete fatty acid oxidation that contributes to muscle insulin resistance (Koves et al., 2008). Thus, the increased *Acaca* response, which is a promoter of fatty acid synthesis, accompanied by higher *Jnk* and *Insr* expression would suggest that leptin deficiency also lead to the development of insulin resistance and glucose intolerance, due a lipid mitochondrial overload by incomplete βoxidation.

According to the changes observed in mitochondrial metabolism and function, the transcription cascade of mitochondrial biogenesis was activated in response to the high-energy demand induced by the absence of leptin. Since defective mitochondrial respiration contributes to mitochondrial proliferation (Marciniak, Marechal et al., 2014), the impaired mitochondrial respiration observed in ob/ob mice probably induces *Ppargc1a* and *Tfam* upregulation as an adaptive response to increase mitochondrial mass and accomplish energy demand. Mitochondrial dynamics are also essential in regulating energy expenditure, maintaining bioenergetics efficiency and determining cell fate. Mitochondrial fusion seems to be particularly important in respiratory active cells since it allows the spreading of metabolites and enzymes while mitochondrial fission contributes to mitophagy or apoptosis (Westermann, 2012). The starvation mode induced by the impairment of satiety response stimulated mitochondrial fusion in order to optimize a defective oxidative phosphorylation and evade cell death-related events.

A shift in energy balance by dietary alterations and increased oxidative stress levels may play an important role in the mechanisms governing protein homeostasis like the UPR (van der Harg et al., 2017, Yilmaz, 2017). In contrast to our findings, several studies have demonstrated increased ER stress and a consequent activation of the UPR in chronic metabolic diseases such as obesity (Hotamisligil, 2010, Ozcan, Cao et al., 2004). Intriguingly, IRE1α/XBP1 and eIF2α pathways were deactivated in muscle fibers from leptin–deficient mice, indicating a reduced accumulation of misfolded proteins in their ER. The only UPR arm that was activated was ATF6α that is implicated in lipid biosynthesis. In fact, a recent study has demonstrated that ATF6α pathway drives lipid biosynthesis, independently of XBP1s (Bommiasamy, Back et al., 2009). In accordance with our results indicating the promotion of lipid anabolism, this leptin deficiency-related ATF6α activation could contribute to the great abundance of cellular lipids. On the other hand, recent findings also relate autophagy to the regulation of ER stress-induced responses and describe a negative feedback in which autophagy negatively regulates the UPR (Senft & Ronai, 2015). Indeed, leptin-deficient mice stimulated autophagy and downregulated protein anabolism by the inhibition of AKT/mTOR pathway that could be critical for UPR deactivation. This may be a consequence of the energy metabolism status acquired by leptin deficiency. The emerging role of hexokinase-II, which is not only implicated in the regulation of metabolism, but also has a market effect on the conserved nutrient-sensing mTOR pathway, emphasizes the importance of understanding selective nutrient adaptation and fuel utilization under conditions of leptin deficiency. A recent study developed in tumor cells revealed that mTOR can relocate to mitochondria, where interacts with hexokinase-II, reprogramming bioenergetics from glycolysis to oxidative phosphorylation, that is required for damage repair and survival (Lu, Qin et al., 2015). However, upon glucose deprivation, hexokinase-II binds to and inhibits mTOR to preserve cellular integrity in neonatal rat ventricular myocytes (Roberts, Tan-Sah et al., 2014). The absence of leptin in obese mice induces a starvation signal and in response to the feeling of limited energy reserves, ob/ob mice increase levels of mitochondrial-bound hexokinase-II, that further support autophagy stimulation thorough mTOR inhibition contributing to the mobilization of diverse cellular energy stores (Kim, Kundu et al., 2011). In addition, this response enhances protein turnover that greatly diminishes the accumulation of missfolded proteins and deactivates IRE1α/XBP1 and eIF2α arms of the UPR. However, this adaptive response may not be based on the removal of dysfunctional mitochondria since ob/ob mice stimulated mitochondrial fusion while suppressing fission, a response that is well established to inactivate mitophagy (Westermann, 2012). Thus, leptin deficiency probably induces an imbalance in mitochondrial homeostasis resulted from an increase in mitochondrial proliferation and a lack of mitophagy process that would explain the accumulation of dysfunctional fused mitochondria and ROS production.

The activation of macroautophagy due the felling of nutrient scarcity stimulates skeletal muscle cells to reutilize their own constituents for energy supply, contributing to skeletal muscle mass reduction. Likewise, this metabolic reprogramming drives the functional fiber-type switch toward the formation of oxidative type I. It was described that PGC-1α that was upregulated in ob/ob mice is also involved in the formation of type I muscle fibers (Bommiasamy et al., 2009). Since glycolytic type II fibers are related to skeletal muscle growth and higher strength (Bommiasamy et al., 2009), the altered energy demand elicited by leptin deficiency stimulates the differentiation of slow fibers and may contribute to the skeletal muscle wasting observed by us. Probably, as a result of a combination of the defective myofiber protein assembly and the fact that slow-twitch muscles fibers contract with lower force than fast ones (Schiaffino & Reggiani, 2011), suppressed excitation-contraction coupling probably occurs in skeletal muscles of leptin-deficient obese mice. This contractile remodeling is probably a consequence of the leptin deficiency-induced metabolic remodeling that is characterized by a redistribution of mitochondrial fuel utilization; so that lipid anabolism is promoted and the high-energy production is stored as fat.

Melatonin and leptin display daily rhythms that may contribute to fuel harvesting and energy homeostasis due the strong interplay between circadian and metabolic systems (Ramsey & Bass, 2011). However, decreased amplitude of nocturnal pineal melatonin peak has been described in obese rodents (Cano, Jimenez-Ortega et al., 2008) and an altered leptin production has also been attributed to a lack of circadian control, having an impact on appetite and energy balance (Mantele, Otway et al., 2012, Nguyen & Wright, 2010). Indeed, melatonin deficiency has been demonstrated to correlate with obesity (Cipolla-Neto, Amaral et al., 2014, Reiter, Tan et al., 2012) and the absence of melatonin leads to leptin resistance (Senft & Ronai, 2015), making leptin treatments ineffective. Then, one can ask whether melatonin supplementation at night normalizes some of the obesity-related metabolic alterations described here. The metabolic normalization observed in melatonin-treated ob/ob mice demonstrates the important role of this neurohormone in maintaining energy homeostasis under conditions of dietary alterations. It has been reported that melatonin improves the utilization of substrates in terms of enhancing the activity of complexes I and IV and increasing ATP production in different pathologies (Martin, Macias et al., 2000, Martin, Macias et al., 2002). A recent publication showed that melatonin treatment also restores the mitochondrial morphology, often devoid of cristae in the heart of obese mice (Stacchiotti, Favero et al., 2017). However, how melatonin counteracts the metabolic alterations resulted from leptin deficiency or leptin resistance remained unknown. Our results demonstrate that melatonin is able to optimize mitochondrial efficiency in the muscle fibers of ob/ob mice enhancing oxidative phosphorylation from complex I and II. Surprisingly, the most relevant finding is that melatonin is able to reduce the exacerbated ATP production in obese animals. Thus, melatonin supplementation under conditions of leptin deficiency not only improves mitochondrial efficiency, but also restores mitochondrial energy demand resembling the requirements of normal-weight mice. As a result of the improvement of mitochondrial function, mitochondrial biogenesis response is attenuated and porin expression suffered a considerable reduction, indicating the recovery of the mitochondrial mass to levels detected in wild-type mice. However, melatonin enhanced *Ppargc1a* that positively affects the expression of ROS-detoxifying enzymes (St-Pierre, Drori et al., 2006), protecting mitochondria from leptin deficiency-induced oxidative stress. Although numerous studies have confirmed the protective effects of melatonin as a potent antioxidant against mitochondrial oxidative stress (Acuna-Castroviejo, Carretero et al., 2012, Ramis, Esteban et al., 2015, Rodriguez, Escames et al., 2007), the readjustment in energy needs in ob/ob treated animals and the improvement of mitochondrial function probably contribute to an adaptation of the redox system. In fact, melatonin reduced lipid peroxidation and increased cytosolic and mitochondrial antioxidant activities. Nevertheless, despite melatonin administration was not able to reduce the 66-kDa isoform of SHC protein activation, mitochondrial catalase activity was much higher, neutralizing the effects mediated by H_2_0_2_ on p66SHC phosphorylation. A recent work has demonstrated that the deletion of p66SHC protects skeletal muscle quality and function against high-fat-induced obesity (Granatiero, Gherardi et al., 2017). Nevertheless, the deficiency of p66SHC in ob/ob mice protect from obesity, but not from glucose intolerance and insulin resistance (Ciciliot, Albiero et al., 2015), suggesting a divergent roles of p66SHC signaling in different eating disorders depending on the bioenergetic status. It is noteworthy that melatonin treatment in ob/ob mice increases the total levels of p66SHC protein, an effect that usually enhance stress-induced apoptosis and cell differentiation (Miyazawa & Tsuji, 2014). Our study demonstrates that melatonin declines the uncontrolled glycolytic capacity and amends the apoptotic response by downregulating the interaction of hexokinase-II with mitochondria. Thus, melatonin could be an important therapeutic against leptin-related disorders due to its capability in modulating antioxidant capability, as well as, enhancing *Pparg* signaling, which is crucial for leptin secretion and upregulating genes involved in glucose uptake, controlling energy homeostasis and improving insulin sensitivity (Ahmadian, Suh et al., 2013). Melatonin has also been able to cope with deficits in oxidative phosphorylation by fuel utilization redistribution and altering mitochondrial dynamics. In contrast to ob/ob mice that show MFN1-mediated mitochondrial fusion, melatonin-treated animals reduced MFN1 levels but presented high MFN2 content. It has been described that the mutation of MFN2 in trophoblast cells lead to the fragmentation of mitochondrial tubules, highlighting the functional importance of MFN2 for embryonic development under conditions of extraordinarily high metabolic rate (Xi, Zhang et al., 2016). Moreover, the reestablishment of fusion capacity by overexpressing MFN2 in mice with severe cachexia attenuates skeletal muscle loss (Chen, Vazquez et al., 2003). IN addition, melatonin has been shown to increases MFN2 response by Notch1 signaling activation preventing myocardial infarction injury (Pei, Du et al., 2016). Therefore, altogether these findings strongly support the therapeutic benefits of melatonin in targeting mitochondrial and lipid metabolism alterations under conditions of the constant exposure to an excessive energy demand induced by the lack of leptin.

These bioenergetic actions of melatonin were accompanied by other metabolic-related effects that favor cell survival and maintenance. It was reported that melatonin is a potent activator of autophagy via mTOR pathway being even capable of counteracting the effects of inhibitors (Choi, Kim et al., 2013). However, although melatonin treatment in ob/ob mice showed further mTOR inhibition, both autophagic flux and p62 were reduced, supporting the previously described protective role of melatonin against protein damage in the liver induced by leptin deficiency (de Luxan-Delgado, Potes et al., 2016). This protective effect of melatonin could be related with its actions in counteracting oxidative damage. Melatonin transforms the exacerbated stress induced by obesity into mild oxidative stress, as indicated by the lower macroautophagy response and higher CMA, whose activation is mediated to counteract mild-oxidative stress (Kiffin, Christian et al., 2004). Consequently, this response contributes to reduce misfolded proteins accumulation, which in turn produces the downregulation of IRE1α/XBP1 and eIF2α pathways. However, ATF6 pathway of UPR has been reestablished by melatonin suggesting a reduction of the lipid anabolism response. Then, melatonin seems to modulate the disturbed nutrient signaling elicited by leptin deficiency, improving proteostasis in skeletal muscle fibers from ob/ob mice by reducing the accumulation of protein aggregates and counteracting ER stress. On the other hand, MFN2 has emerged as a linker of mitochondrial function, ER stress and insulin signaling being essential for glucose homeostasis. The lack of MFN2 triggers hydrogen peroxide production, ER stress, and active JNK (Munoz, Ivanova et al., 2013, Sebastian, Hernandez-Alvarez et al., 2012). Then, the upward regulation of MFN2 induced here by melatonin treatment in ob/ob mice further support that this hormone play a crucial role in the modulation of all these pathways improving skeletal muscle function.

In summary, melatonin has potential musculoskeletal benefits under the pleiotropic effects of leptin mutation, modifying fuel utilization and targeting the switch between oxidative type I fibers to glycolytic type II fibers. Therefore, melatonin could be a potential therapeutic agent for leptin deficiency-induced obesity and leptin-related disorders by partly mimicking leptin signaling in several physiological functions and by regulating mitochondrial bioenergetics.

## Materials and methods

### Animals

Sixteen six male wild-type (C57BL/6J) mice and sixteen six male leptin-deficient ob/ob (B6.VLepob/J) mice at six weeks of age were purchased from Charles River Laboratory (Charles River Laboratories, SA, Barcelona, Spain). All mice were maintained on a 12:12 h dark/light cycle at 22 ± 2°C with tap water and standard chow diet *ad libitum*. This study has developed twice, one for general studies and another one for cytosolic and mitochondrial isolated experiments. The Oviedo University Animal Care and Use Committee approved the experimental protocol. All in vivo studies were carried out according to the Spanish Government Guide and the European Community Guide for Animal Care (Council Directive 86/609/EEC).

### Treatment

All experimental animals were subjected to two weeks acclimatization period under standard laboratory conditions before treatment. Wild-type (W) and ob/ob (Ob) mice were randomized into four experimental groups of eight animals each: non-treated control groups (WC and ObC, respectively) and melatonin-treated groups (WM and ObM, respectively). Two h after lights off (ZT14), melatonin (Sigma Aldrich, St Louis, MO, USA) diluted in the minimum volume of ethanol (0.5%) was subcutaneously injected daily at a dose of 500μg/kg body weight for four weeks. Non-treated animals received a comparable dose of the vehicle.

Animals were sacrificed by decapitation, and the skeletal muscle from the hind limb of each mouse was removed. Muscle samples were washed with saline solution and immediately frozen in liquid nitrogen and stored at 80°C until further use. If not indicated otherwise, muscle (0,4 g) from each mouse were homogenized using an Ultra-turrax homogenizer (Ultra-Turrax T25 digital; IKA, Staufen, Germany) in 2 mL of lysis buffer (50 mM phosphate buffer, pH 7.5, 1 mM NaF, 1 mM Na3VO4, 1 mM PMSF, 0.1% Triton-X 100) and centrifuged for 6 min at 1500 g and 4°C. Supernatants were collected and Bradford method (Bradford, 1976) was used to measure protein content.

### Macroscopic parameters

All animals were weighed at baseline and at the end of the experiment. At the sacrifice, body composition was assessed by the quantification of muscle and fat mass weights, the length of lower limb perimeter and the height of each mouse. BMI, SMI, FFMI and L-ASMI were calculated in the four experimental groups to characterize the sarcopenic muscle. Food intake was measured per cage twice a week.

### Histological analysis

Pieces of skeletal muscle from the hind limb were fixed by immersion in 4% formaldehyde (20910.294, MERCK, Darmstadt, Germany), embedded in paraffin by standard methods and cut into 5-7μm thick sections. Sections were stained by periodic acid-Schiff (PAS) to evaluate muscle fibers type pattern. In each skeletal muscle piece, four different levels separated by 300μm were collected, and in each level, two random high-power fields (HPF) were analyzed using a NIKON Eclipse E200 microscope (Nikon Corp., Tokyo, Japan) (the size of the HPF for this microscope is 0.196 mm^2^). The proportion of type-II fibers stained darker than type I fibers were quantified. Quantification was done by observers who were blinded to the genotype of the animals and treatments.

### Mitochondria isolation

Mitochondrial isolation from skeletal muscle was performed following García-Cazarin and colleagues protocol. (Garcia-Cazarin, Snider et al., 2011) The tissue was washed with PBS, and immediately cut slowly in little pieces to been incubated in PBS solution with 10mM EDTA and 0.01% of trypsin on ice. After 30 minutes incubation period, muscle pieces were transferred to mitochondrial isolation buffer 1 (10mM EDTA, 215mM D-mannitol, 77mM sucrose, 20mM HEPES, 0.1% BSA (fatty acid free) pH 7.4), homogenized using a Potter-Elvehjem homogenizer and centrifuged for 10 min at 700g and 4°C. The supernatant containing cytosolic and mitochondrial fractions was transferred to a new refrigerated centrifuge tube and centrifuged again at 10500g for 10 min at 4°C. The resulting supernatant that contained cytosolic fraction was then collected. The pellet, corresponding to mitochondrial fraction was gently resuspended in mitochondrial isolation buffer 2 (6mM EGTA, 215mM D-mannitol, 77mM sucrose, 20mM HEPES, 0.1% BSA (fatty acid free) pH 7.4) and centrifuged at 10500g for 10 min at 4°C. Finally, supernatant was discarded and the mitochondrial isolated pellet was resuspended in mitochondrial isolation buffer 2. Protein content in cytosolic and mitochondrial extracts were quantified by the Bradford method. (Bradford, 1976)

### Oxidative stress status

Lipid peroxidation (LPO) was studied by the measure of reactive aldehyde malondialdehyde (MDA) and 4-hydroxy-2-(E)-nonenal (4-HNE) end-products. The content of MDA and 4-HNE were determined in skeletal muscle homogenates using a LPO assay kit from Calbiochem (No. 437634, San Diego, CA, USA) based on the condensation of the chromogene 1-methyl-2-phenylindole with either MDA or 4-HNE. (Gerard-Monnier, Erdelmeier et al., 1998) Cytosolic SOD and mitochondrial SOD activities (EC 1.15.1.1), which catalyze the dismutation of the superoxide anion (O_2.-_) to hydrogen peroxide (H_2_O_2_), were measured based on the inhibition of hematoxylin autoxidation to the colored compound hematein, according to the method developed by Martin and colleagues. (Martin, Dailey et al., 1987) CAT activity was assayed in cytosolic and mitochondrial fractions using the method reported by Lubinsky and Bewley,(Lubinsky & Bewley, 1979) which is based on the breakdown of H_2_O_2_ into O_2_ and H_2_O. GSH-Px catalyzes the oxidation of reduced glutathione (GSH) to oxidized glutathione (GSSG) by the reduction of H_2_0_2_ to H_2_0. The GSSG produced is immediately reduced to GSH by nitocinamide adenine dinucleotide phosphate (NADPH) and glutathione reductase (GR) action. GSH-Px and GR were measure in cytosolic and mitochondrial samples and the rate of NADPH consumption was monitored to determined both activities according to Kum-Tatt and colleagues. (Kum-Tatt, Tan et al., 1975)

### Oxygen consumption

Mitochondrial oxygen consumption was determined using a Clark-type oxygen electrode (Oxygraph, Hansatech Instruments Ltd, Pentney, UK) following the protocol developed by Silva and Oliveira. (Silva & Oliveira, 2012) 500 μg of mitochondrial suspension was introduced into the chamber in 1mL of mitochondrial respiration buffer (5mM MgCl_2_,215mM of D-mannitol, 6.5μM KH_2_PO_4_, 20μM EGTA, 15mM sucrose, 4mM HEPES, 0.1% BSA (fatty acid free) pH 7.4). Complex I and complex II respirations were measured by the addition of specific substrates (complex I: 10mM glutamate + 5mM malate, 175nmol ADP, 1μg oligomycin, 1μM carbonyl cyanide-4-(trifluoromethoxy) phenylhydrazone (FCCP); complex II: 5mM succinate + 1μM rotenone, 125nmol ADP, 1μg oligomycin, 1μM FCCP). RCR was calculated as the ratio between maximal O_2_ consumption stimulated by ADP (state 3) and the respiration in absence of ADP or without ATP synthesis (state 4), that determines the coupling between substrate oxidation and phosphorylation. (Silva & Oliveira, 2012) ADP/O was also obtained as the amount of ADP added per amount of O_2_ consumed during state 3. (Silva & Oliveira, 2012)

### ATP measurement

The Adenosine 5’-triphosphate (ATP) Bioluminescent Assay kit (FLAA, Sigma-Aldrich) was used to determine ATP content in cytosolic and mitochondrial fractions. This assay measures the light emission with a luminometer based on the ATP consumption when firefly luciferase catalyzes D-luciferin oxidation.

### Mitochondrial transmembrane electric potential

Isolated mitochondria (2.5 mg/mL) were incubated with 16.7 μM red-fluorescing MitoTracker Red FM (M22425, Thermo Fisher Scientific, Waltham, MA, USA) for 20 minutes at 30°C. A volume of 8 μL of stained mitochondria was analyzed using a Leica TCS SP2 AOBS Laser Scanning Spectral Confocal Microscope (Leica Biosystems, Wetzlar, Germany) to determine the mitochondria membrane potential. Fluorescence per mitochondrial mass (area) was quantified with Image J.

### Enzyme-linked immunosorbent assay (ELISA)

TNF-α (KMC3011, Life Technologies, Waltham, MA, USA) and IL-6 (KMC0061, Life Technologies, Waltham, MA, USA) ELISA kits were used to assess the inflammatory response in skeletal muscle tissue. All assays were performed following manufacturers’ protocols.

### Peptide mass fingerprinting

Aliquots of muscle homogenate (25 μg of protein per sample) were solubilized in Laemmli sample buffer (BioRad Laboratories, Inc., CA, USA) and boiled at 100°C for 5 min to load onto SDS-PAGE gels. Molecular weight rang marker composed by a mixture of blue-stained recombinant proteins (Precision Plus Protein All Blue Standars, BioRad) were also loaded onto the gels to identify proteins molecular weights. Coomassie Brilliant Blue R-250 dye (BioRad) was used to stain one-dimensional gels for protein detection. Gel images were semiquantitatively analyzed using Image Studio Lite 3.1.4 software for Macintosh (LI-COR Biosciences, NE, USA).

Protein bands of interest were manually excised from Coomassie stained gels and submitted for peptide mass fingerprint identification at the Inbiotec S.L. (León, Spain) proteomics laboratory. Samples were processed and analyzed with a 4800 Proteomics Analyzer matrix-assited laser desorption ionization time-of-flight (MALDI-TOF/TOF) mass spectrometer (ABSciex, MA, USA) according to the previously described methods of Oliván et al. (Oliván, Fernández-Suárez et al., 2016) A database search on Mascot Generic Files combining MS and MS/MS spectra was performed using Mascot v 2.2 from Matrix Science through the Global Protein Server v 3.6 (ABSciex). When the Mascot score was greater than 85 points, the identified protein was considered a valid candidate.

### Western blotting

Muscle tissue homogenates, cytosolic or mitochondrial extracts (50-100 μg of protein per sample) were denaturalized at 100°C for 5 min in a Laemmli buffer (BioRad) to be separated by electrophoresis in SDS-PAGE gel at 200 V and transferred to a polyvinylidene fluoride membranes (Immobilon TM-P; Millipore Corp., MA, USA) at 350 mA. After blocking membranes for 1 h at room temperature with 10% (w/v) skim milk in TBS (50mM Tris-HCl, (pH 7.5) and 150mM NaCl), membranes were incubated overnight at 4°C with the respective primary antibodies: AIF (sc-13116; Santa Cruz Biotechnology); ATF6α (sc-22799, Santa Cruz Biotechnology, Texas, USA); BAX (2772, Cell Signaling); Beclin-1 (4445, Cell signaling, Danvers, MA); CaMKII (3362, Cell Signaling); CHOP (L63F7) (2895, Cell Signaling); CI-20 (NDUFB8) (ab110242, Abcam, Cambridge, UK); CII-30 (SDHB) (ab14714, Abcam); CIII-Core II (UQCRC2) (ab14745, Abcam); CIV-I (MTCO1) (ab14705, Abcam); CV-a (ATP5A) (ab14748, Abcam); cyclophilin D (ab110324 (MSA04), Abcam); cytochrome C (ab110252, Abcam); DRP1 (D6C7) (8570S, Cell Signaling); eIF2α (5324, Cell Signaling); Hexokinase-II (2867; Cell Signaling); IRE1α (3294, Cell Signaling); LC3 (PD014, Medical & Biological Laboratories Co., LDT, Japan); MFN1 (sc-50330; Santa Cruz Biotechnology); MFN2 (D2D10) (9482S, Cell Signaling); MURF1 (ab77577; Abcam); p62 (H00008878-M01, Abnova, Taipe, Taiwan); p70S6K (9202; Cell Signaling); phospho-AKT (9271, Cell Signaling); phospho-CaMKII (3361, Cell Signaling); phospho-eIF2α (3398, Cell Signaling); phospho-mTOR (5536; Cell Signaling); phospho-p70S6K (9206; Cell Signaling); phospho-RYR1 (ab59225, Abcam); phospho-SHC/p66 (566807; Calbiochem); PI3K (4255; Cell Signaling); porin (MSA03) (ab14734, Abcam); SHC (610879; BD Trasduction Laboratories); XBP1(sc-8015; Santa Cruz Biotechnology). Each antibody was previously diluted in blocking buffer (2% (w/v) skim milk in TBS). After three 10 min washes in TBS-T (TBS containing 0.05% Tween-20), membranes were incubated with the corresponding horseradish peroxidase-conjugated secondary antibody (Sigma-Aldrich, Misuri, USA) for 1 h at room temperature. After three 10 min washes in TBS-T, membranes were developed using a chemiluminesent horseradish peroxidase substrate (WBKLS0500, Millipore Corp., Darmstadt, Germany) according to manufacturer’s protocol. Densities of proteins bands were analyzed quantitatively with Image Studio Lite 3.1.4 software (LI-COR Biosciences, Nebraska, USA). Variations in the levels of the typical housekeeping proteins (GAPDH, β-actin and α-tubulin) were found, so Ponceau S staining was used to ensure equal protein loading. (Fortes, Marzuca-Nassr et al., 2016)

### RNA extraction and RT-qPCR analysis

RNA was isolated from mouse s keletal muscle tissue using TRI reagent (T9424, Sigma Aldrich). Total RNA extraction levels were quantified using NANO DROP 2000 (Thermo Fisher Scientific). High capacity cDNA reverse transcription kit (4368814, Applied Biosystems, Foster City, CA, USA) was used to synthesize complementary DNA (cDNA) from total RNA extracts following the manufacture’s instructions. Expression levels of genes were determined in StepOne real-time PCR system (Applied Biosystems) by quantitative real-time PCR (RT-qPCR) reactions using Power SYBR Green PCR Master Mix (4367659; Applied Biosystems) and the specific pairs of primers: *Acaca*, forward: 5’-AATGGCATTGCAGCAGTGAA-3’, reverse: 5’-CACATAGTGATCTGCCATCTTAATGTATT-3’; *Adipor1*, forward: 5’-CCCACCATGCCATGGAGA-3’, reverse: 5’-GCCATGTAGCAGGTAGTCGTTGT-3’; *Adipor2*, forward: 5’-CAGGAAGATGAGGGCTTTATGG-3’, reverse: 5’-GAGGAAGTCATTATCCTTGAGCCA-3’; *Atp2a1*, forward: 5’-CTGACCGCAAGTCAGTGCAA-3’, reverse: 5’-GGATGGACTGGTCAACCCG-3’; *Calstabin1*, forward: 5’-GGGGATGCTTGAAGATGGAA-3’, reverse: 5’-TTGGCTCTCTGACCCACACTC-3’; *Gapdh*, forward: 5’-CAATGACCCCTTCATTGACC-3’, reverse: 5’-TGGAAGATGGTGATGGGATT-3’; *Insr*, forward: 5’-TGAACGCCAAGAAGTTTGTG-3’, reverse: 5’-CAGCCAGGCTAGTGATTTCC-3’; *Jnk*, forward: 5’-GATTGGAGATTCTACATTCACAG-3’, reverse: 5’-CTTGGCATGAGTCTGATTCTGAA-3’; *Lamp2a*, forward: 5’-GAAGTTCTTATATGTGCAACAAAGAGCAG-3’, reverse: 5’-CTAAAATTGCTCATATCCAGCATGATG-3’; *Ppargc1a*, forward: 5’-GACTTGGATACAGACAGCTTTCTGG-3’, reverse: 5’-GCTAGCAAGTTTGCCTCATTCTCT-3’; *Ppara*, forward: 5’-TGAAGAACTTCAACATGAACAAG-3’, reverse: 5’-TTGGCCACCAGCGTCTTC-3’; *Pparg*, forward: 5’-ACTATGGAGTTCATGCTTGTGAAGGA-3’, reverse: 5’-TTCAGCTGGTCGATATCACTGGAG-3’; *Ryr1*, forward: 5’-AAGGCGAAGACGAGGTCCA-3’, reverse: 5’-TTCTGCGCGTTGCTGTGG-3’; *Tfam*, forward: 5’-ACCTCGTTCAGCATATAACGTTTATGTA-3’, reverse: 5’-GCTCTTCCCAAGACTTCATTTCAT-3’; *Xbp1*, forward: 5’-GAGGAGAAGGCGCTGAGGA-3’, reverse: 5’-CCTCTTCAGCAACCAGGGC-3’. Thermocycling conditions were as follows: a holding stage for 10 min at 95°C; then a cycling stage of 40 15 s cycles at 95°C and 1 min at 60°C; and finally a 15 s melt curve stage at 95°C, 1 min at 60°C and 15 min at 95°C. The average cycle threshold (Ct) value at which each gene was detectable was calculated. The Ct of the GAPDH was used for normalization. Relative changes in gene expression levels were determined using the 2^-ΔΔCt^ method. (Livak & Schmittgen, 2001)

### Statistical analysis

The statistical software package SPSS 20.0.0 (SPSS Inc., Chicago, IL, USA) and GraphPad Prism 6.0 (GraphPad Software, Inc. CA, USA) for Macintosh were used for all statistical analyses and graph design. Data are mean values ± standard deviation of the mean (SD). The normality of the data was analyzed using the Kolmogorov-Smirnov test or the Mann-Whitney non-parametric test for continuous variables. The effect of genotype and melatonin treatment was determined by ANOVA followed by Bonferroni *post-hoc* test. A p<0.05 was considered statistically significant.

## Acknowledgements

This work was supported by the Instituto de Salud Carlos III (Spanish Ministry of Economy and Competitiveness) (PI13/02741 and PI17/02009); the PCTI (Principado de Asturias) (GRUPIN-071); and FEDER funds. J.C. B-M. is a pre-doctoral fellow at the Instituto de Investigación Sanitaria del Principado de Asturias (IISPA). Y.P. is a FISS pre-doctoral fellow at the Instituto de Salud Carlos III (Spanish Ministry of Economy and Competitiveness) (FI14/00405). We are members of the INEUROPA and CIBERFES networks.

## Author contributions

Y.P. performed the experiments and analyzed the data. A. D-L, J.C. B-M and Z.P-M. helped in data acquisition and analysis. B. de L-D. and A. R-G. helped in performing mitochondrial oxygen consumption experiments. I.M-V. helped in performing oxidative stress experiments. J. G-R. and J.J.S. helped in histological analysis. B. C. provided expertise in statistics. B.C., I.V-N. and A. C-M. conceptualized, designed and supervised the study. I.V-N. and A.C-M. provided suggestions and revised the manuscript. Y.P. wrote the manuscript.

